## Supplementary Information for "Heterogeneous efflux pump expression underpins phenotypic resistance to antimicrobial peptides"

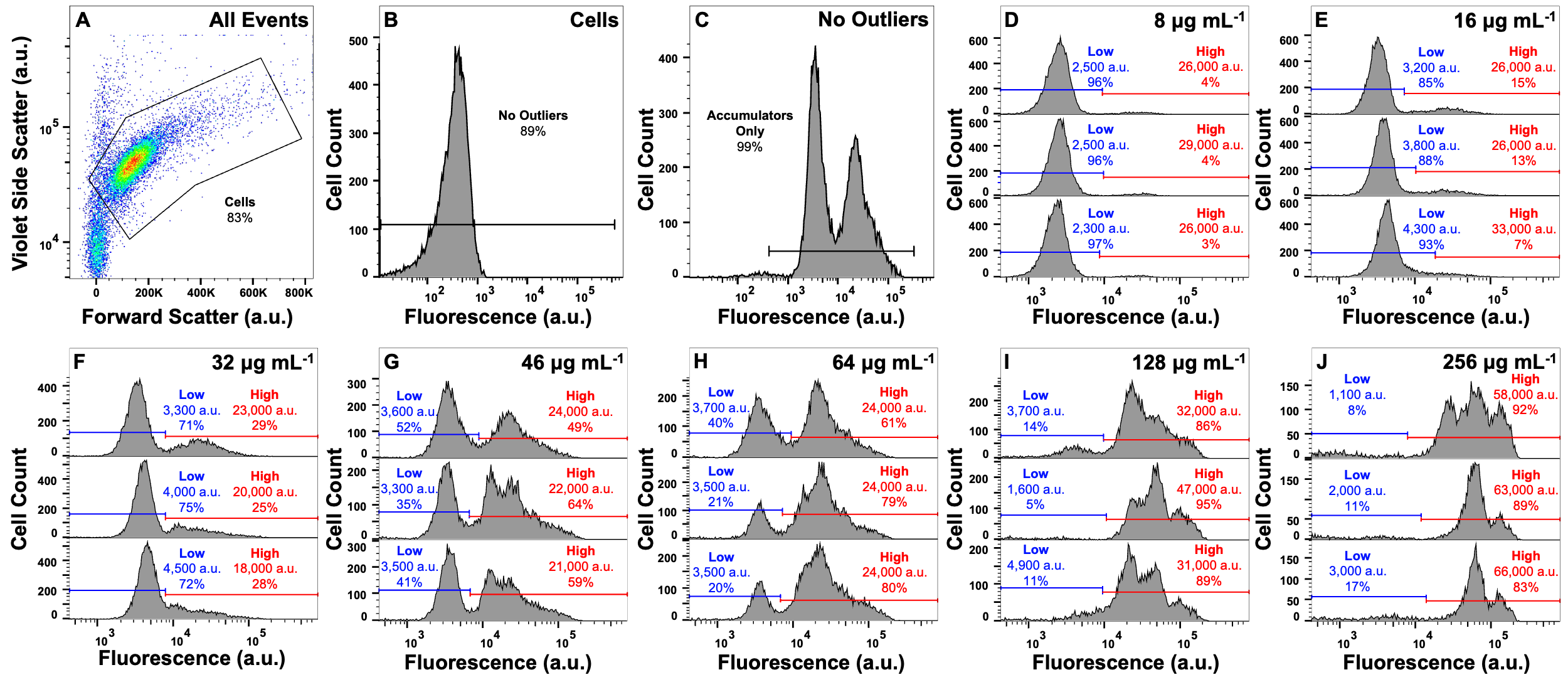

**Figure S1.** **Gating strategy and dose-dependent response to tachyplesin-NBD treatment.** (A-C) Gating strategy for all flow cytometric assays. Bacteria were gated and separated from cellular debris using forward scatter and violet side scatter (A). Background noise was then further separated based on cellular autofluorescence measured on the FITC-A channel for cells not treated with fluorescent-peptides (B). An additional gate on the FITC-A channel was then used for cells treated with fluorescent-peptides to further separate background noise (C). (D-J) Median fluorescence and proportion of low (blue) and high (red) tachyplesin-NBD accumulators within *E. coli* BW25113 stationary phase populations treated with 8 μg mL^-1^ (3.2 μM) (D), 16 μg mL^-1^ (6.3 μM) (E), 32 μg mL^-1^ (12.7 μM) (F), 46 μg mL^-1^ (18.2 μM) (G), 64 μg mL^-1^ (25.4 μM) (H), 128 μg mL^-1^ (50.7 μM) (I), or 256 μg mL^-1^ (101.5 μM) (J) tachyplesin-NBD in M9 at 37 °C for 60 min. Each of the three graphs in each panel show data for 10,000 events collected from an independent biological replicate. The horizontal lines in each graph represent the manual gating applied.

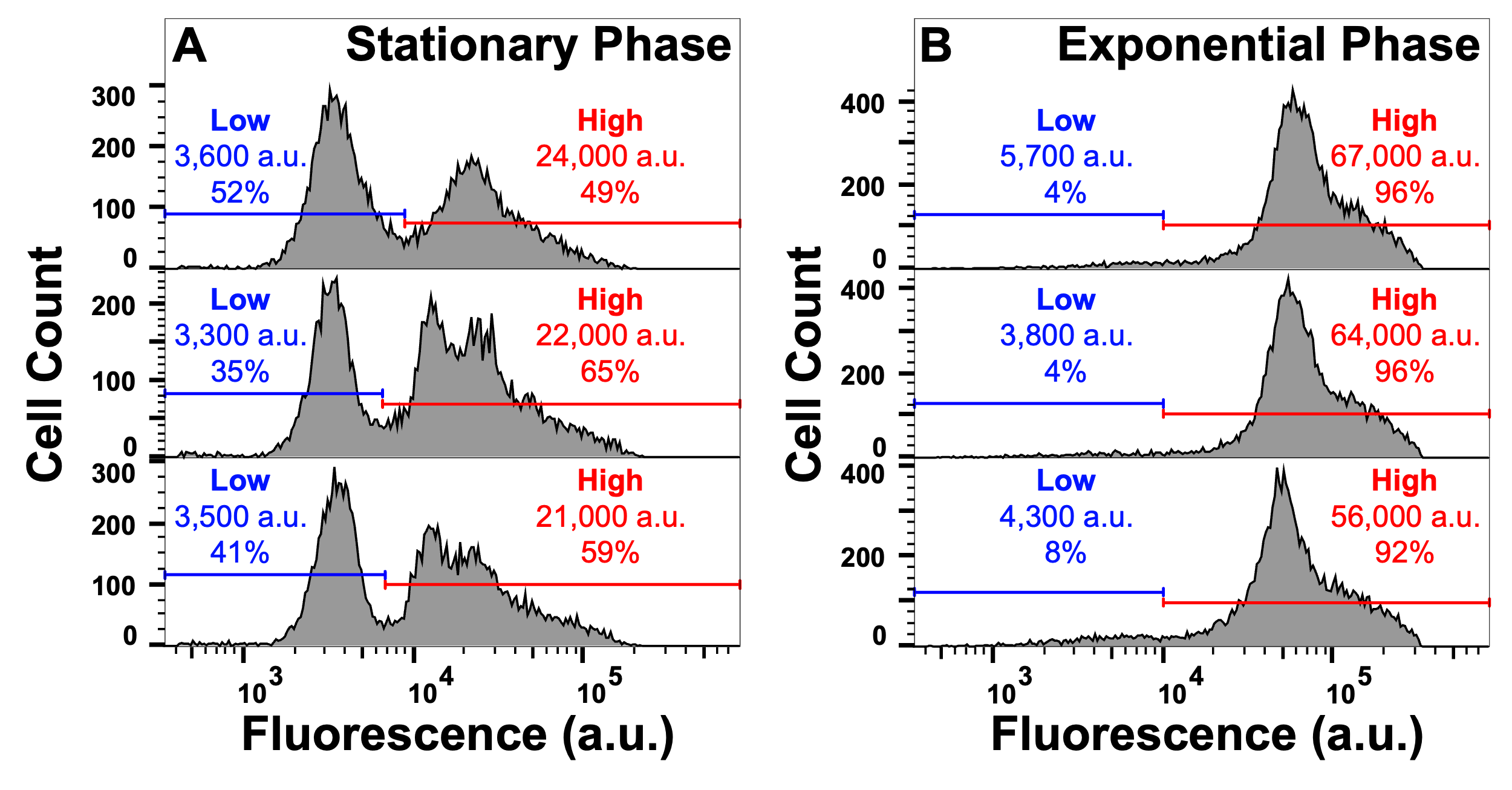

**Figure S2. Tachyplesin-NBD accumulation in stationary and exponential phase *E. coli* BW25113.** Stationary (A) and exponential (B) phase *E. coli* BW25113 were treated with 46 μg mL^-1^ (18.2 μM) tachyplesin-NBD in M9 at 37 °C for 60 min and single-cell fluorescence measured via flow cytometry. The blue and red horizontal lines in each graph represent the gating applied and the median fluorescence and proportion of cells in the corresponding gate are reported within each graph. Each of the three graphs in each panel show data collected from an independent biological replicate.

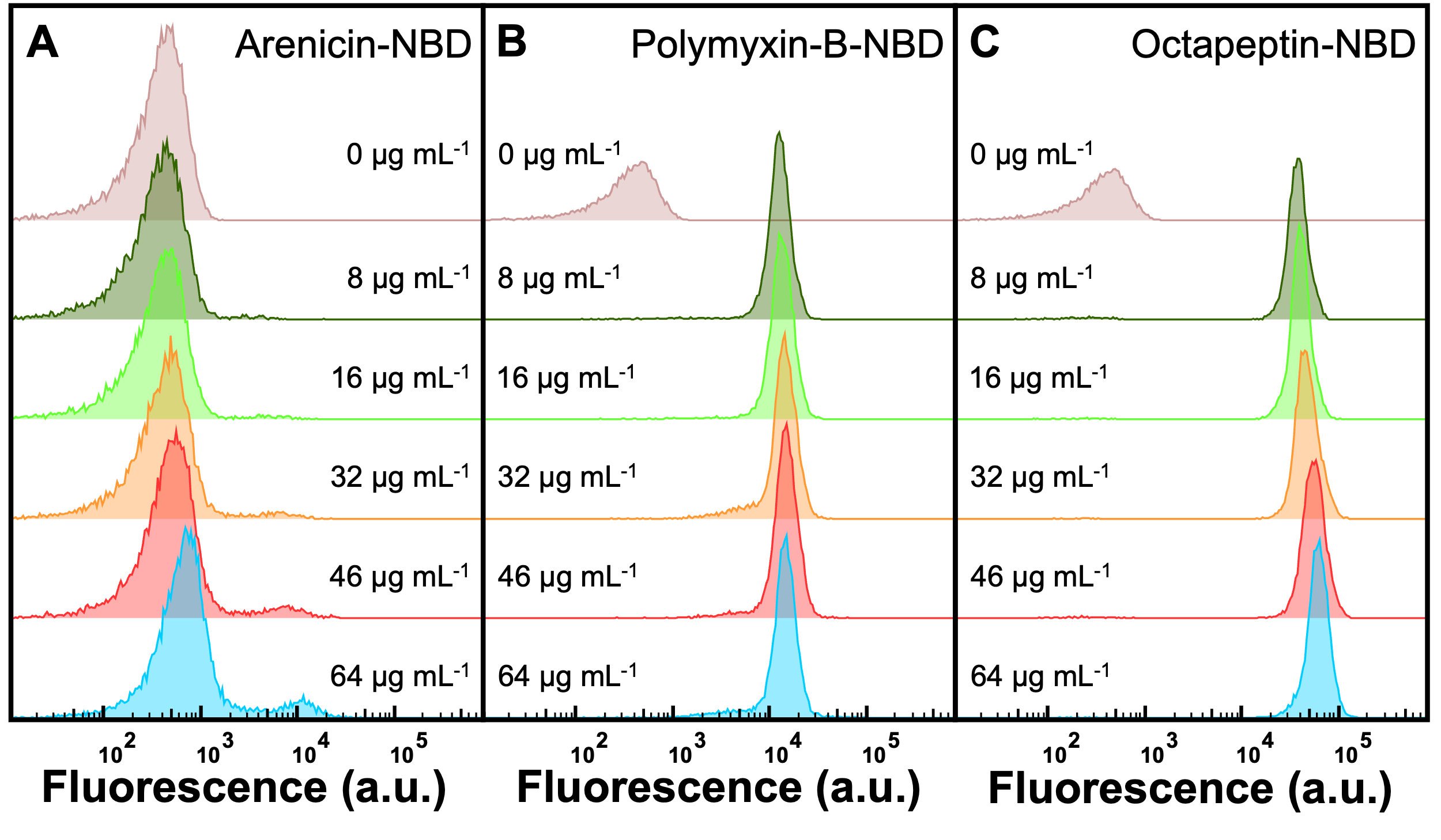

**Figure S3. Dose-dependent response to arenicin-NBD, polymyxin-B-NBD, and octapeptin-NBD treatment.** (A-C) Stationary phase *E. coli* BW25113 was treated with 0, 8, 16, 32, 46, or 64 μg mL^-1^ of arenicin-NBD (0.0, 2.8, 5.7, 11.3, 16.3, 22.7, 45.4, or 90.7 μM) (A), polymyxin-B-NBD (0.0, 7.0, 13.9, 27.9, 40.0, 55.7, 111.4, or 222.8 μM) (B), and octapeptin-NBD (0.0, 6.1, 12.3, 24.5, 35.3, 49.1, 98.2, or 196.3 μM) (C) in M9 at 37 °C for 60 min. Data are representative of data collected from three biological replicates.

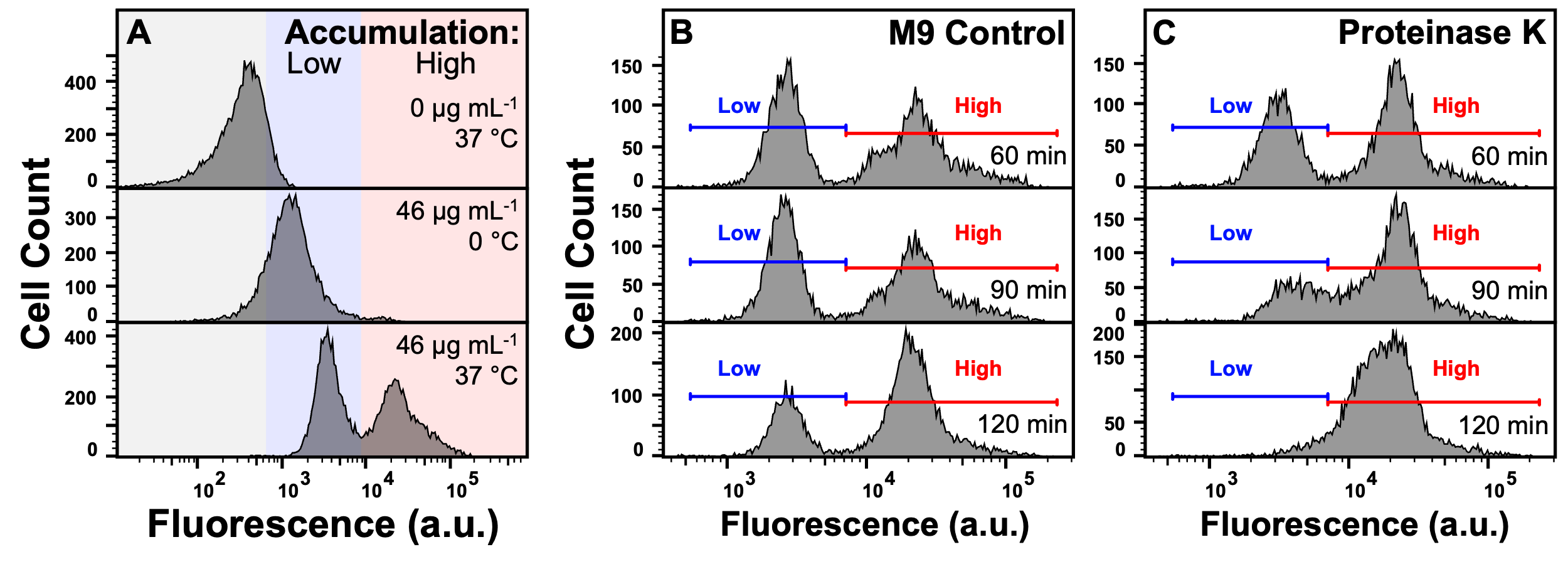

**Figure S4. Impact of environmental temperature and proteinase K on tachyplesin-NBD accumulation.** (A) Stationary phase E. coli BW25113 were treated with 0 or 46 μg mL^-1^ (0 or 18.2 μM) tachyplesin-NBD in M9 at 37 °C or on ice for 60 min. Each panel shows data representative of three independent biological replicates. (B and C) Stationary phase E. coli BW25113 were treated with 46 μg mL^-1^ (18.2 μM) tachyplesin-NBD in M9 at 37 °C for 60 min. Cells were washed and then incubated in either M9 (B) or 20 μg mL^-1^ (0.7 μΜ) proteinase K in M9 (C) at 37 °C over 120 min. Each panel shows data representative of three independent biological replicates.

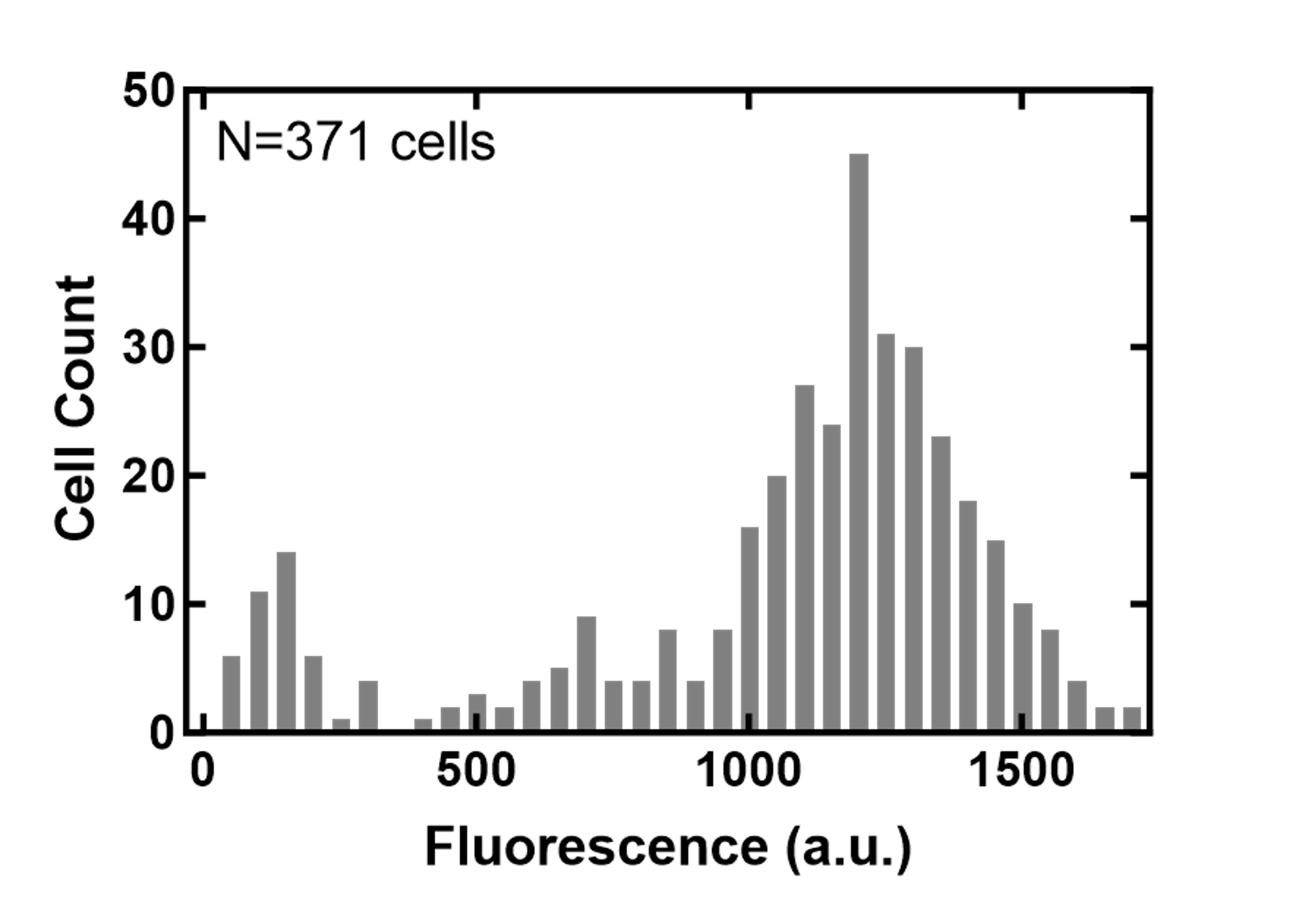

**Figure S5. Distribution of single-cell tachyplesin-NBD accumulation in the microfluidic mother machine.** Distribution of tachyplesin-NBD accumulation in stationary phase *E. coli* BW25113 treated with 46 μg mL^-1^ (18.2 μM) tachyplesin-NBD in M9 at 37 °C for 60 min then washed with M9 at 37 °C for 60 min in the microfluidic mother machine.

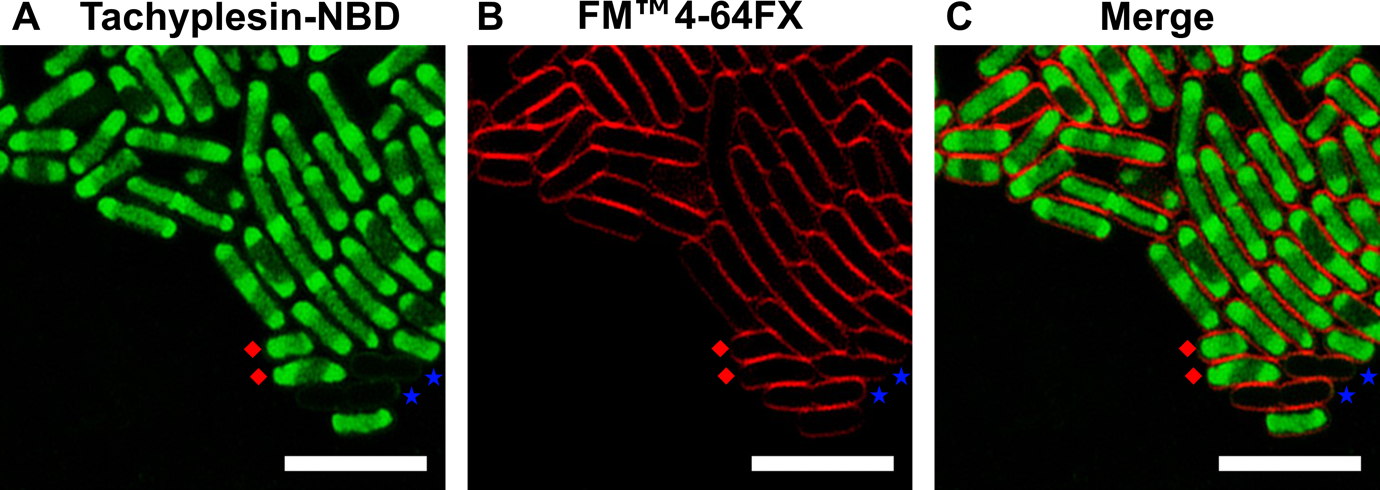

**Figure S6. Representative live-cell confocal microscopy images of E. coli treated with tachyplesin-NBD and FM™ 4-64FX.** Exponential phase E. coli ATCC 25922 were treated with 16 μg mL^-1^ (6.3 μM) tachyplesin-NBD at 37 °C for 30 minutes, followed by staining with 5 μg mL^-1^ (6.3 μM) FM™ 4-64FX, a membrane-specific dye on ice. The samples were then visualised using confocal microscopy. (A) Image showing tachyplesin-NBD fluorescence (green), indicating AMP accumulation in the cells. (B) Image showing FM™ 4-64FX fluorescence (red), highlighting the bacterial cell membrane. (C) Merged images from panels A and B, showing the intracellular accumulation of tachyplesin-NBD. Blue stars and red diamonds indicate representative low and high accumulators, respectively. Scale bars represent 5 μm. Data was collected from three independent biological replicates.

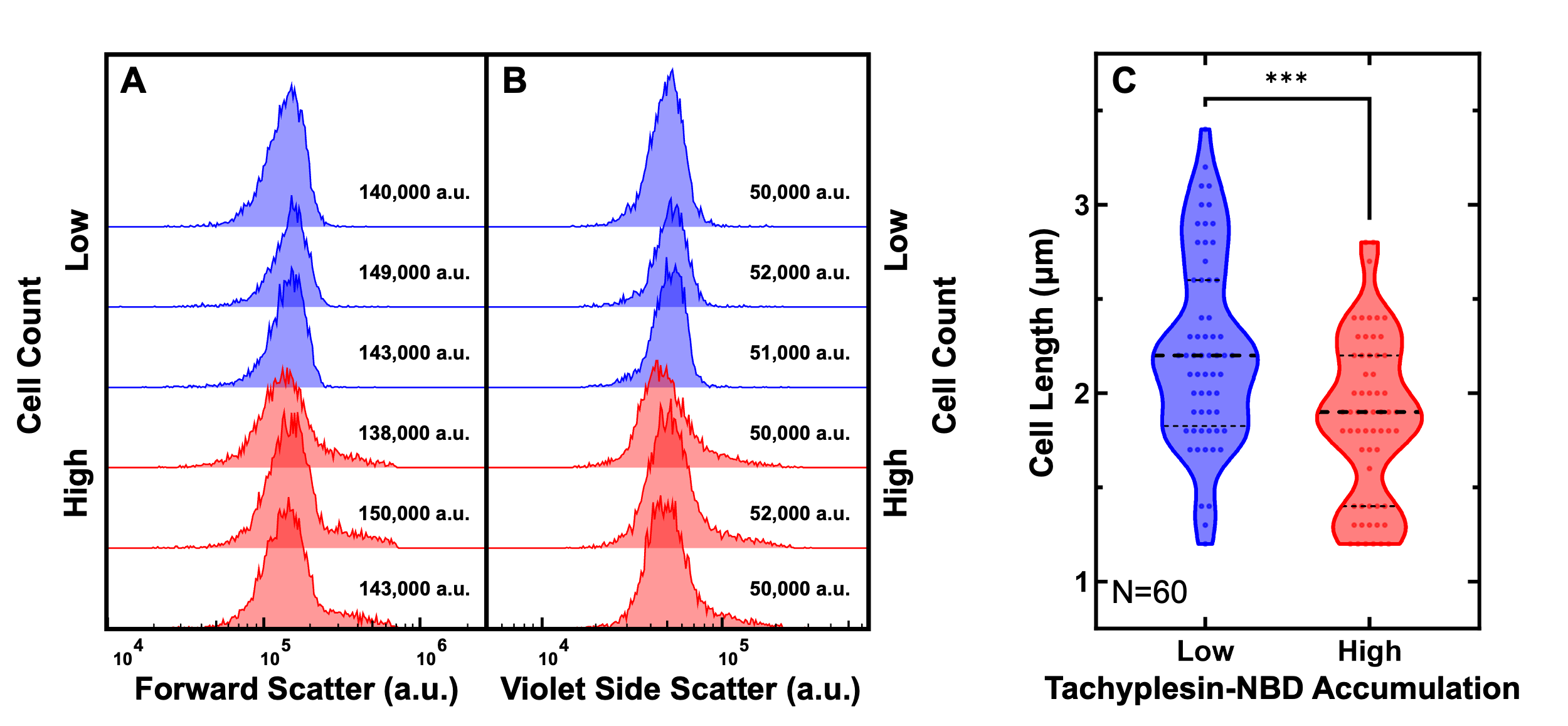

**Figure S7. Comparison of cell sizes between low and high tachyplesin-NBD accumulators.** (A and B) Distribution of forward scatter (A) and violet side scatter (B) of low (blue) and high (red) accumulators after treatment with 46 μg mL^-1^ (18.2 μM) tachyplesin-NBD in M9 at 37 °C for 60 min. Corresponding medians reported in each graph. Each histogram in each panel show data collected from an independent biological replicate. (C) Distribution of cell lengths of low and high tachyplesin-NBD accumulators measured by using the microfluidics-based microscopy platform. Stationary phase *E. coli* was treated with 46 μg mL^-1^ (18.2 μM) tachyplesin-NBD in M9 at 37 °C for 60 min then washed with M9 at 37 °C for 60 min in the microfluidic mother machine. Classification of low and high tachyplesin-NBD accumulators was further validated by propidium iodide staining (see Figure 2G). Statistical significance was assessed using an unpaired two-tailed nonparametric Mann-Whitney U test. P-value: *** p=0.0002. Data was collected from three independent biological replicates.

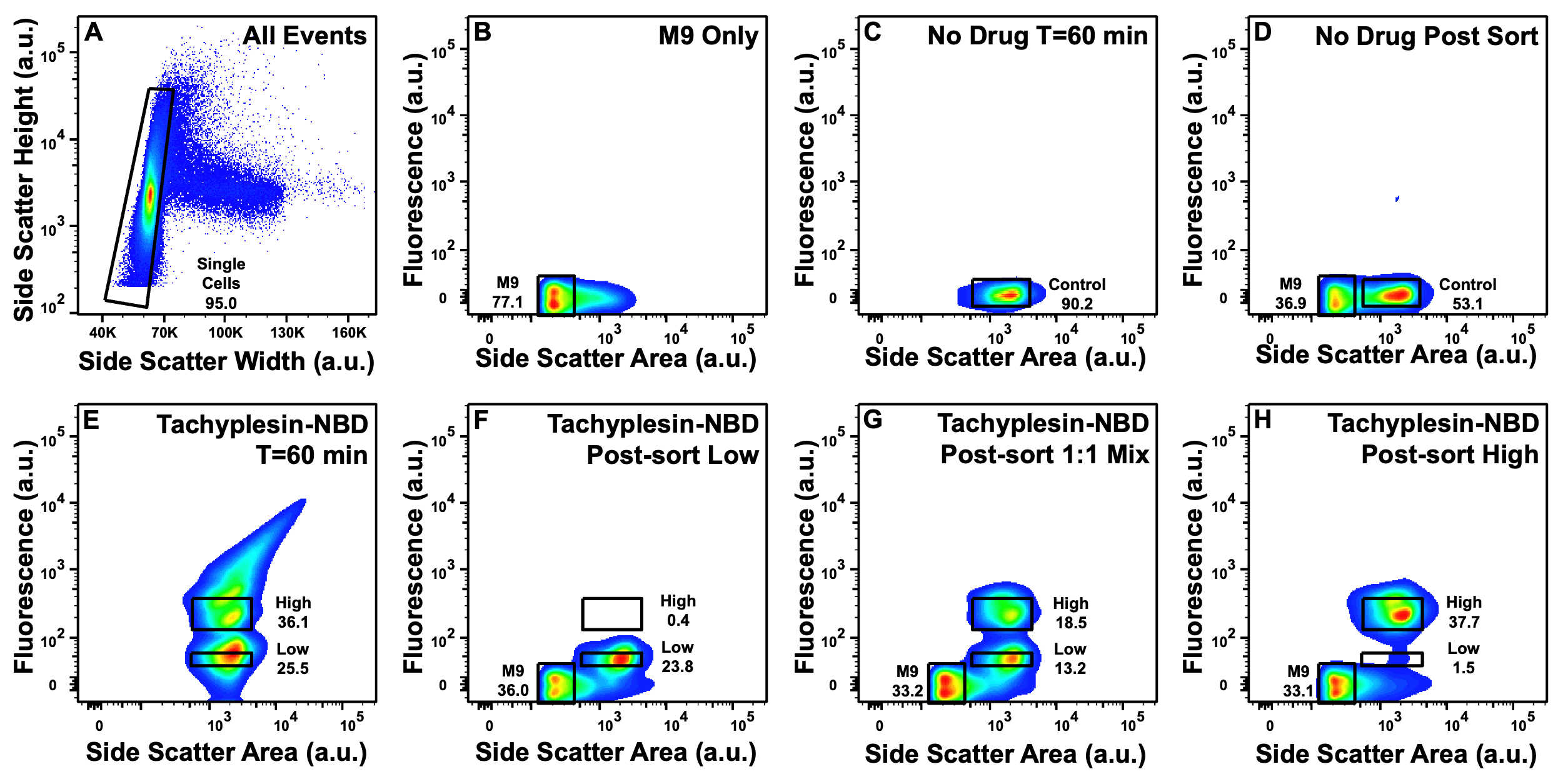

**Figure S8. Fluorescence-activated cell sorting of untreated cells, low and high tachyplesin-NBD accumulators.** (A) Gating on side scatter width against side scatter height was applied on all events to separate single cells from cell aggregates. (B-H) Representative plots of fluorescence and side scatter of: M9 only (B); individual *E. coli* incubated in M9 at 37 °C for 60 min (C); post-sort analysis of *E. coli* sorted through control gate in C (D); individual *E. coli* incubated in 46 μg mL^-1^ (18.2 μM) tachyplesin-NBD at 37 °C for 60 min (E); post-sort analysis of *E. coli* sorted through the low gate in E (F); post-sort analysis of a 1:1 mixture of *E. coli* sorted through the low and high gates in E (G); post-sort analysis of *E. coli* sorted through the high gate in E (H). Data was collected from four independent biological replicates.

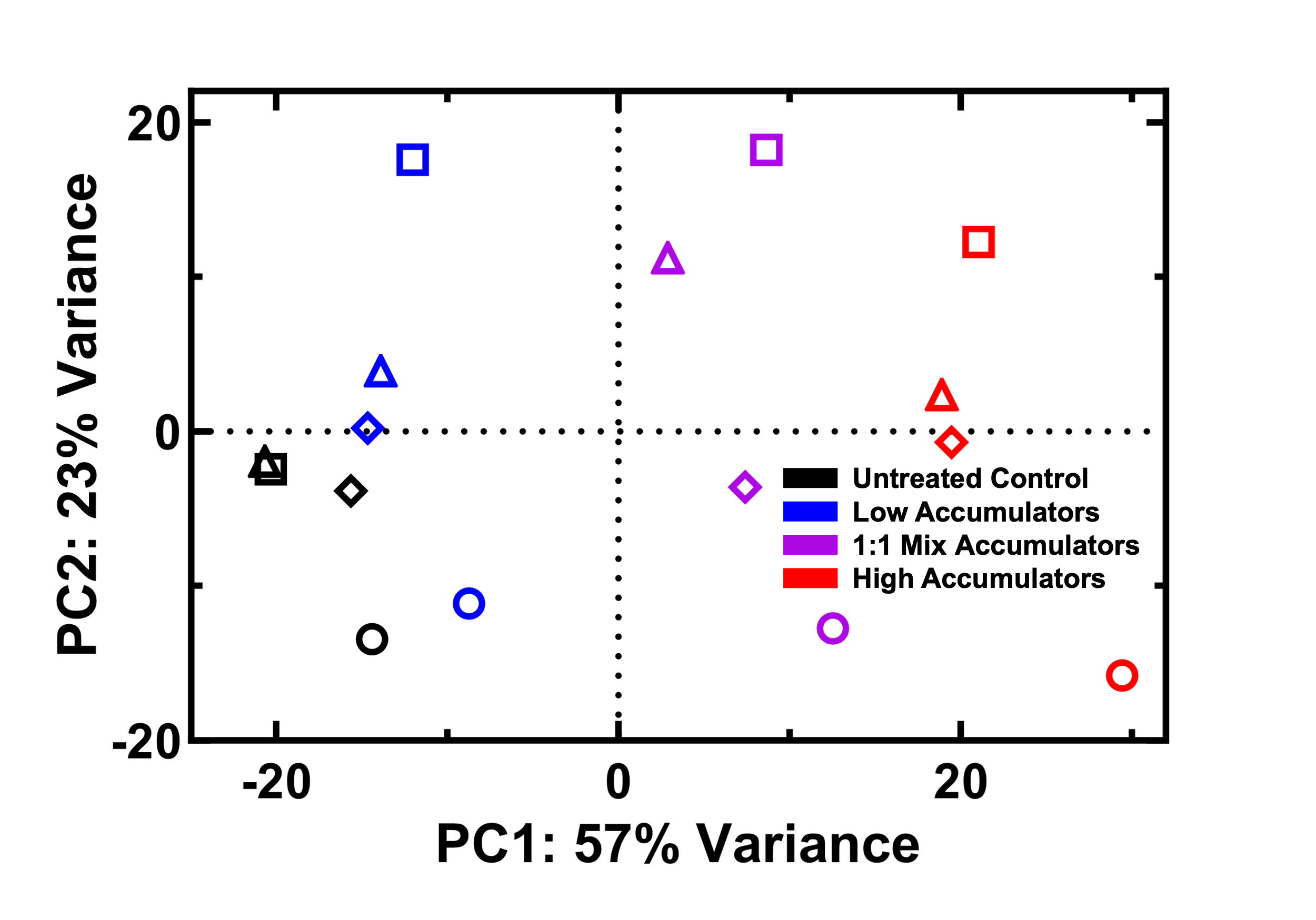

**Figure S9. Principal component analysis of transcriptomes.** Cells were analysed after 60 min treatment in M9 only (black) or 46 μg mL^-1^ (18.2 μM) tachyplesin-NBD, then sorted for low (blue) and high (red) accumulators, and 1:1 mix (purple) accumulators (generated by a 1:1 mixture of low and high accumulators). Circles, squares, diamonds, and triangles indicate four separate biological replicates.

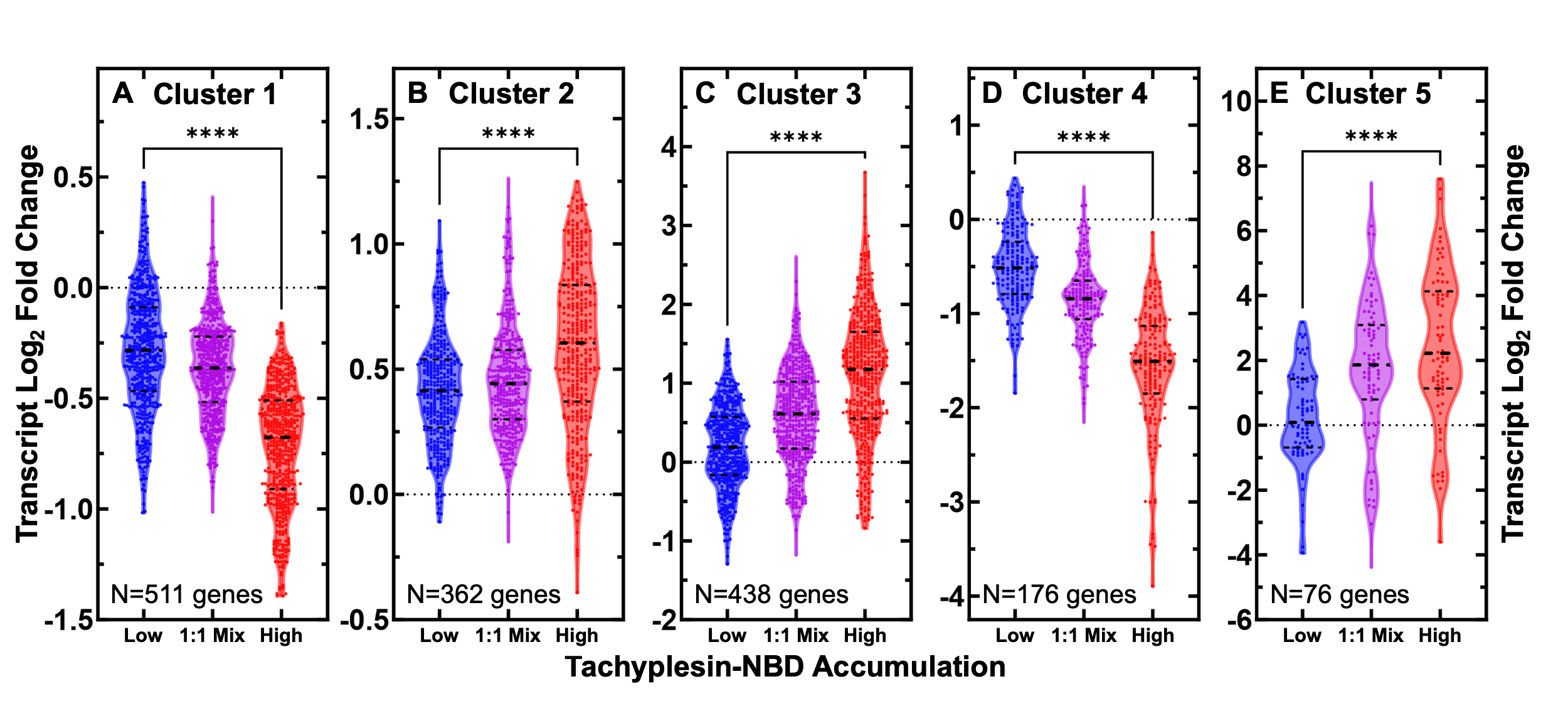

**Figure S10. Clustering of differentially regulated genes in low, 1:1 mix, and high accumulators of tachyplesin-NBD.** (A-E) Log_2_ fold changes in transcript reads of genes in cluster 1 (A), 2 (B), 3 (C), 4 (D) and 5 (E) in low (blue), 1:1 mix (purple), and high (red) tachyplesin-NBD accumulators relative to the control treatment. Each point represents a single gene, dashed lines indicate the median and quartiles of each distribution. Dotted lines represent a log_2_ fold change of 0. Statistical significance was tested using a paired Wilcoxon nonparametric test (due to non-normally distributed data) with a two-tailed p-value and confidence level at 95%. **** p<0.0001. The full list of genes belonging to each cluster are reported in Data Set S2. Data was collected from four independent biological replicates.

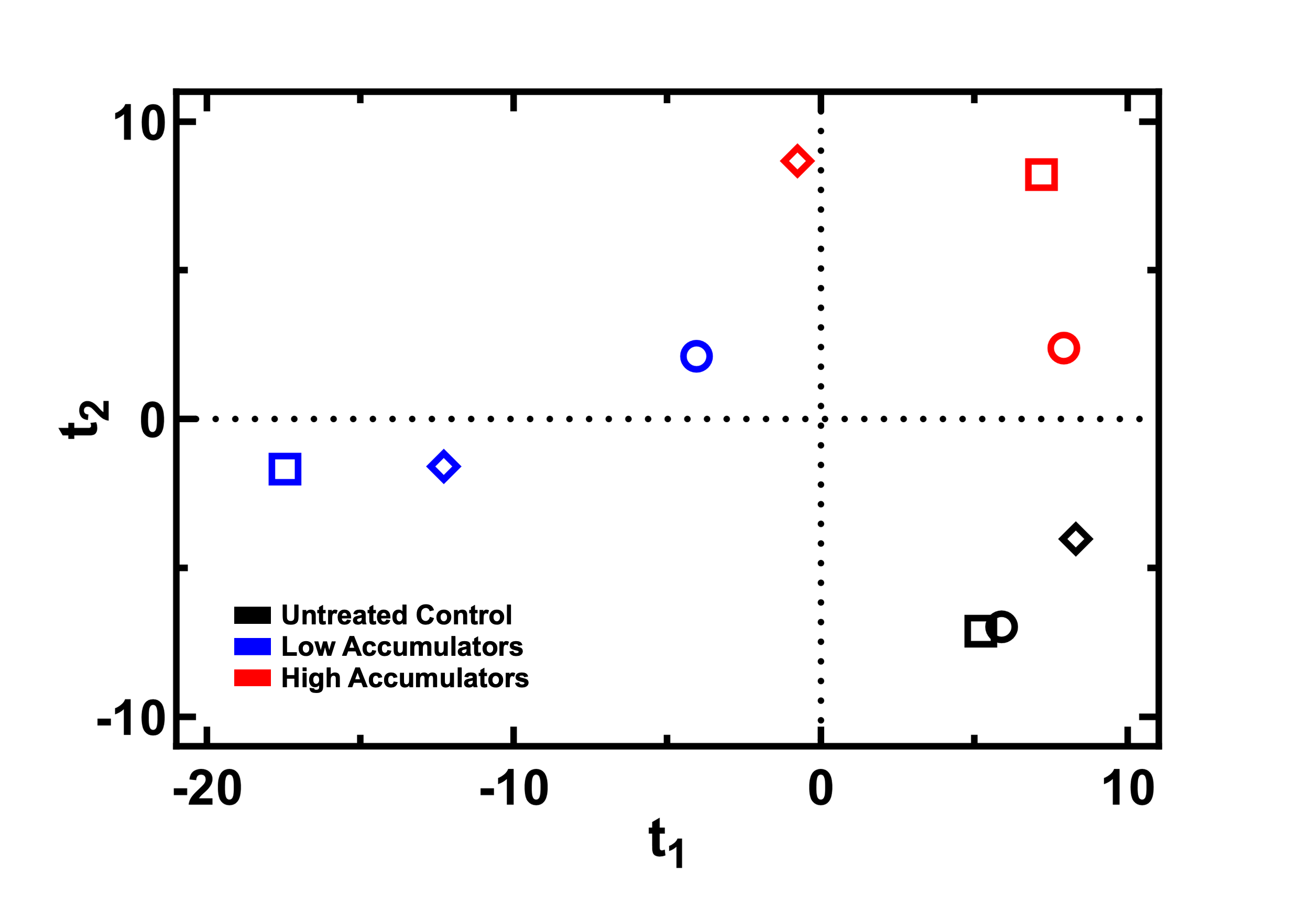

**Figure S11. Clustering of lipidomes of low and high accumulators of tachyplesin-NBD.** Orthogonal partial least squares-discriminant analysis of the lipidomes measured for bacteria treated in M9 only (black) or sorted low (blue) or high (red) accumulators after 46 μg mL^-1^ (18.2 μM) tachyplesin-NBD treatment for 60 min. Circles, squares, and diamonds indicate three separate biological replicates.

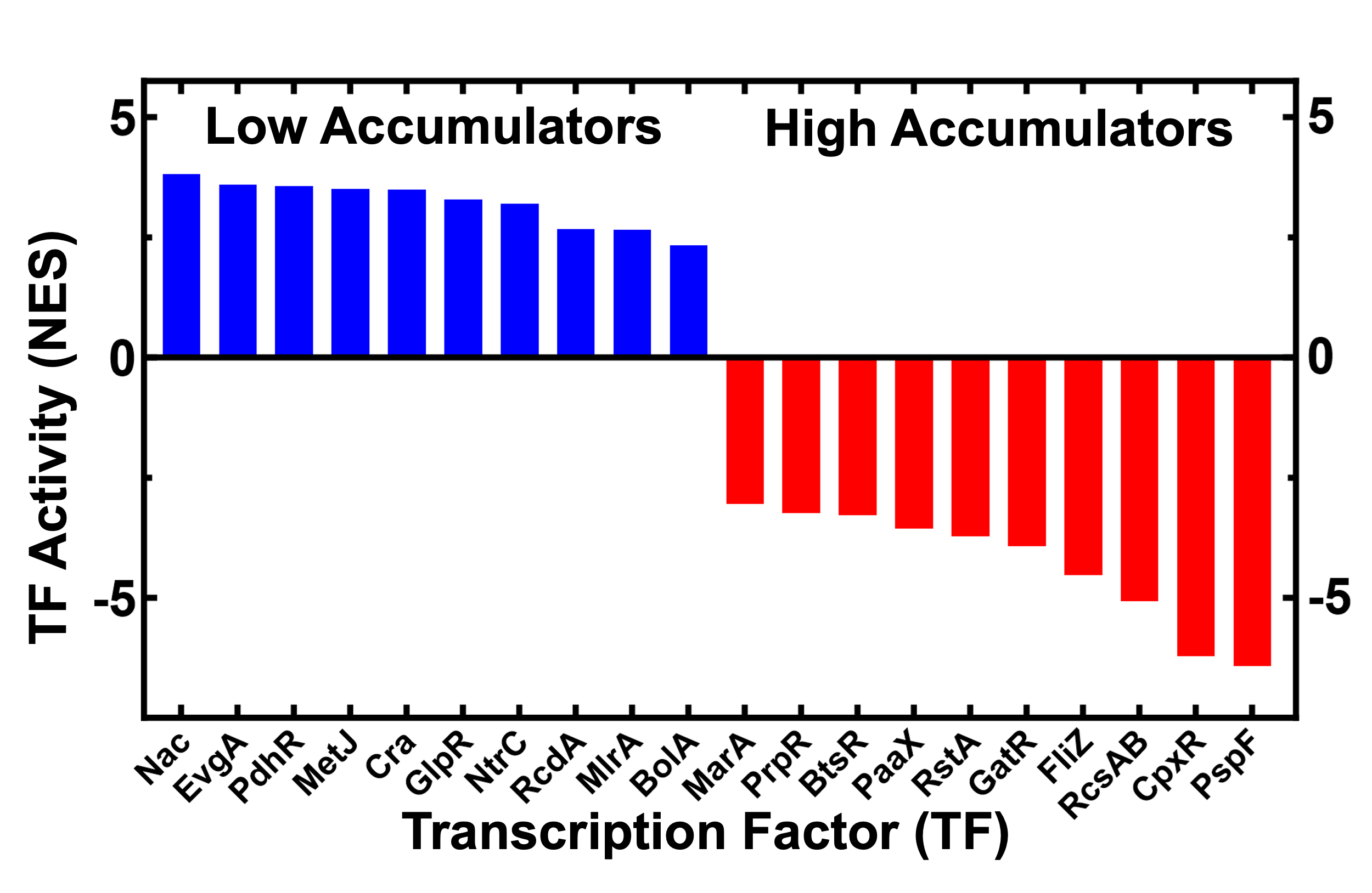

**Figure S12. Inferred transcription factor (TF) activity for the ten TFs with highest inferred activity in low and high tachyplesin-NBD accumulators.** TF activity reported as normalised enrichment scores (NES), for low and high accumulators (blue and red bars, respectively). Activity was inferred using Data Set S2 and a full list of TFs and the genes they regulate within each cluster is reported in Data Set S4.

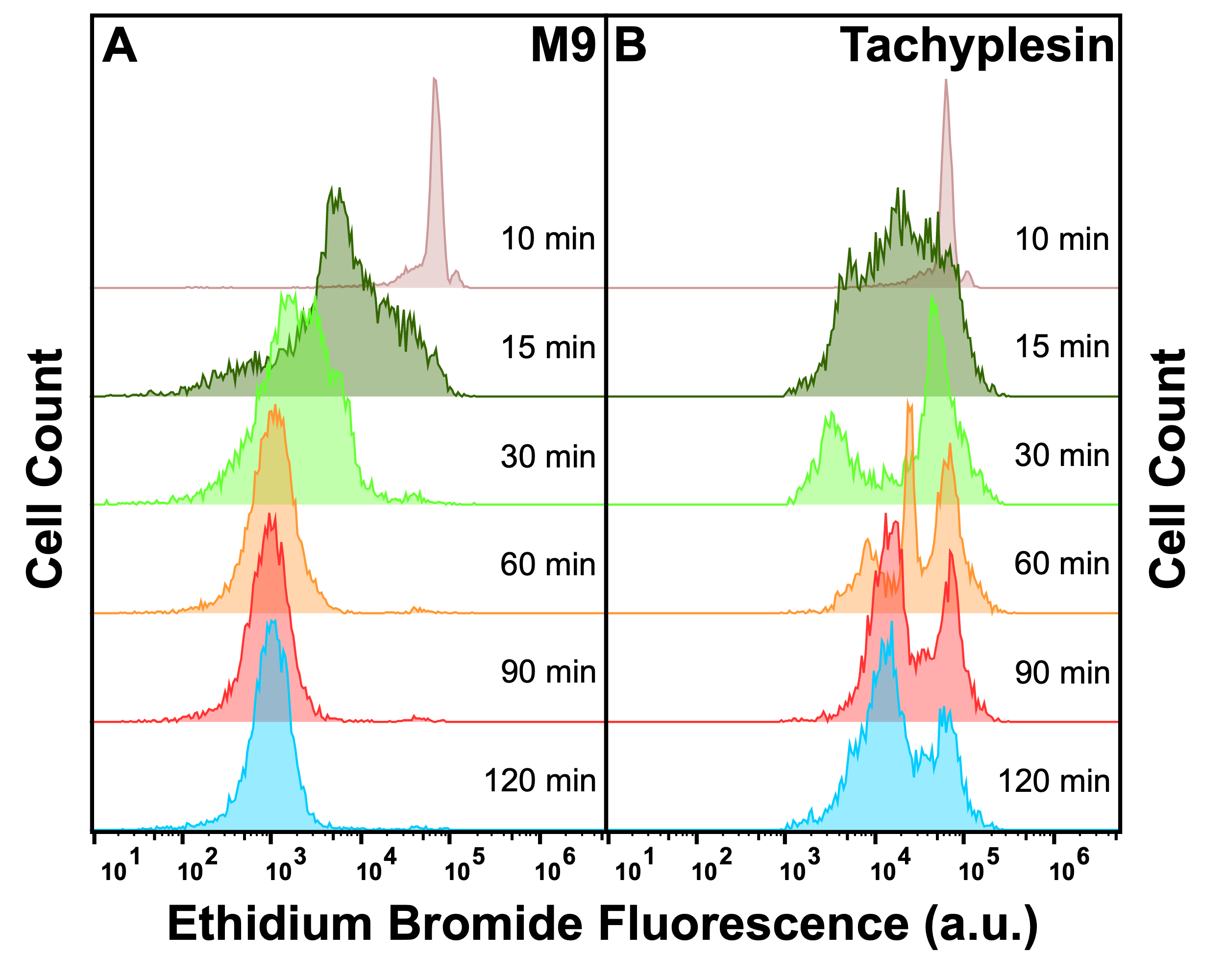

**Figure S13. Ethidium bromide efflux in the presence and absence of unlabelled tachyplesin-1.** E. coli cells were pre-loaded with ethidium bromide (EtBr) by adding 100 μg mL^-1^ (254 μM) EtBr to a stationary phase culture 90 min before reaching 17 h incubation at 37 °C and 200 rpm. Cells were then washed to remove extracellular EtBr before resuspending in either M9 (A) or 46 μg mL^-1^ (20.3 μM) unlabelled tachyplesin. Single-cell EtBr fluorescence was then measured via flow cytometry over 120 min. Histograms are representative of three independent biological replicates.

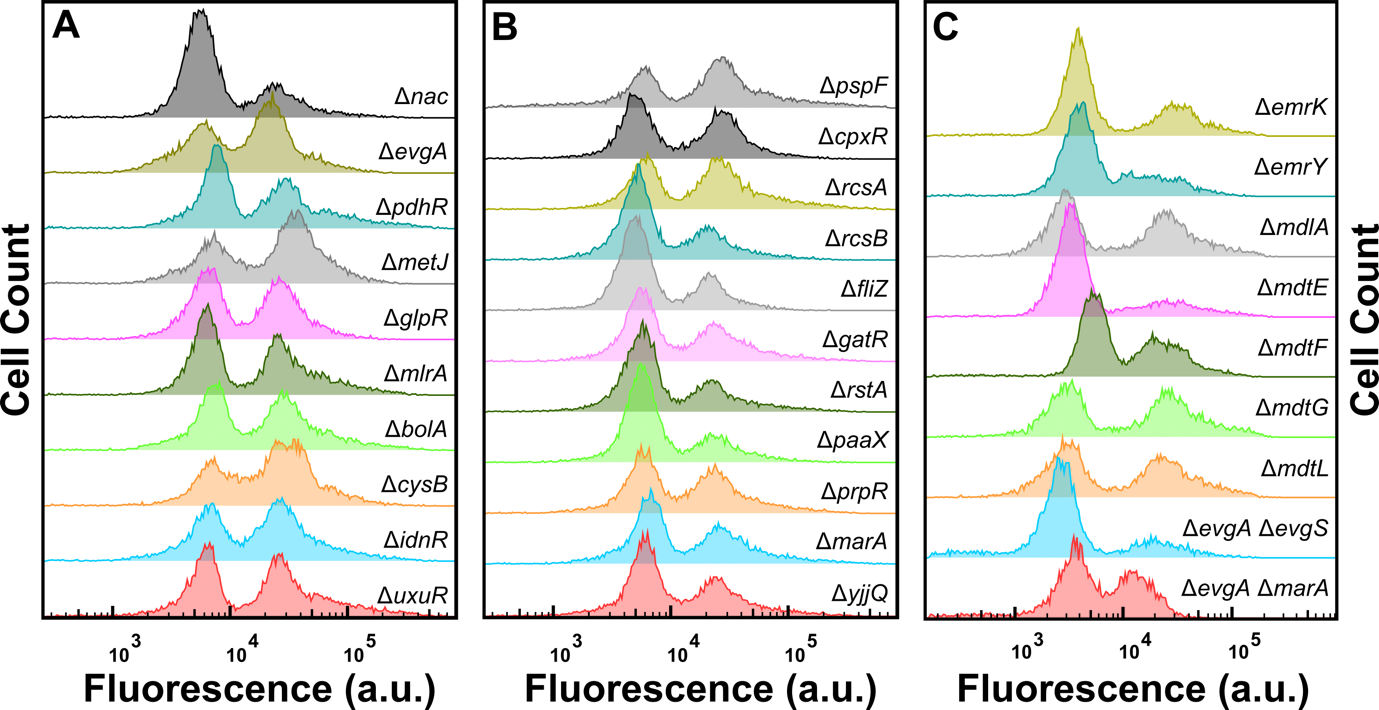

**Figure S14. Impact of efflux component and transcription factor deletion on tachyplesin-NBD accumulation.** Tachyplesin-NBD accumulation in E. coli BW25113 single or double gene-deletion mutants lacking either efflux components or regulators of efflux components or the transcription factors reported in Figure S12. Stationary phase populations of each mutant were treated with 46 μg mL^-1^ (18.2 μM) tachyplesin-NBD at 37 °C for 60 min and single-cell fluorescence was measured via flow cytometry. Histograms are representative of three independent biological replicates.

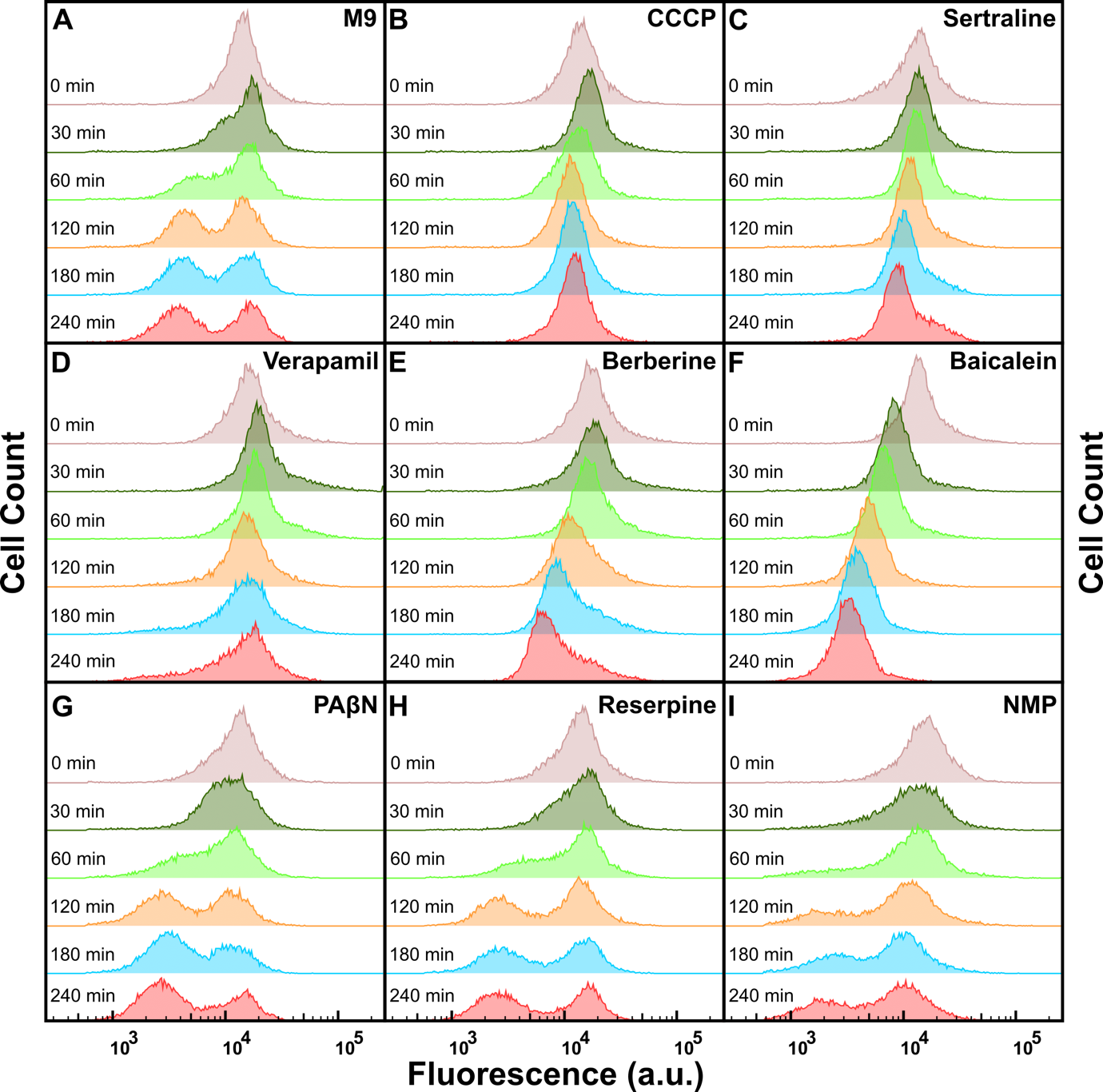

**Figure S15. Tachyplesin-NBD efflux in the presence of different efflux pump inhibitors.** (A-I) Temporal dependence of the distribution of tachyplesin-NBD accumulation in stationary phase E. coli treated in 46 μg mL^-1^ (18.2 μM) tachyplesin-NBD in M9 for 15 min, then washed to remove extracellular tachyplesin-NBD and transferred into M9 (A), CCCP (50 μg mL^-1^ or 244 μM) (B), sertraline (30 μg mL^-1^ or 98 μM) (C), verapamil (50 μg mL^-1^ or 110 μM) (D), berberine (250 μg mL^-1^ or 743 μΜ) (E), baicalein (25 μg mL^-1^ or 93 μM) (F), phenylalanine-arginine beta-naphthylamide (PAβN; 20 μg mL^-1^ or 53 μM) (G), reserpine (20 μg mL^-1^ or 33 μM) (H) and 1-(1-naphthylmethyl)piperazine (NMP; 100 μg mL^-1^ or 442 μM) (I). Each distribution is representative of three independent biological replicates.

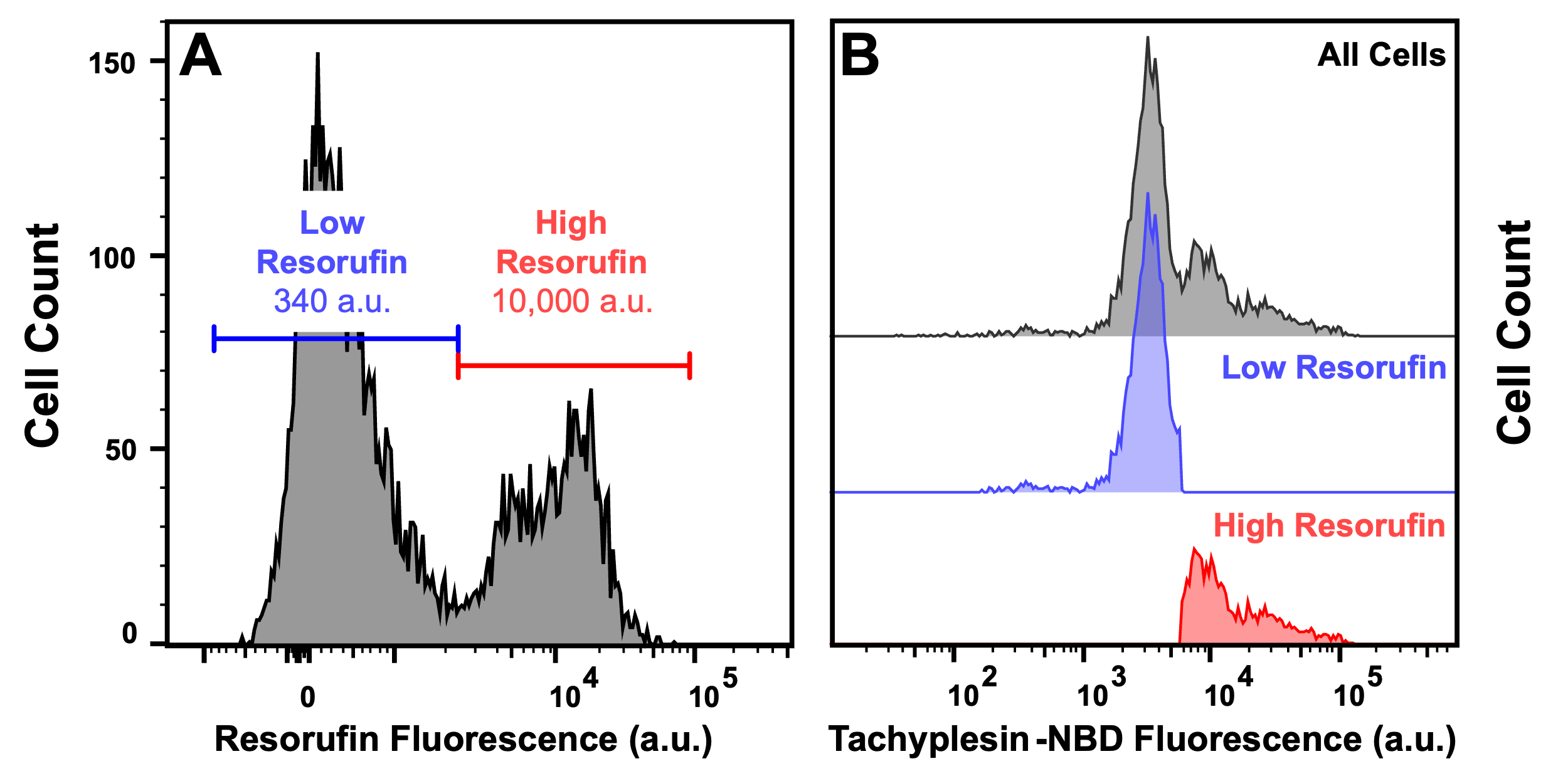

**Figure S16. Differences in metabolic activity of low and high tachyplesin-NBD accumulators.** Stationary phase E. coli BW25113 were treated with 46 μg mL^-1^ (18.2 μM) tachyplesin-NBD in carbon-free M9 at 37 °C for 60 min. Cells were washed and then incubated in 1 μM resazurin and 50 μM CCCP in carbon-free M9 at 37 °C for 15 min. Single-cell fluorescence of resorufin and tachyplesin-NBD fluorescence was measured simultaneously with flow cytometry. (A) Single-cell resorufin fluorescence in all cells. The blue (low) and red (high) horizontal lines in each graph represent the gating applied and the median resorufin fluorescence of cells in the corresponding gate is reported within each gate. (B) Single-cell tachyplesin-NBD fluorescence measured in all cells (top panel); cells measured with low resorufin fluorescence (middle panel, showing cells only within the low resorufin gate in panel A); cells measured with high resorufin fluorescence (bottom panel, showing cells only within the high resorufin gate in panel A).

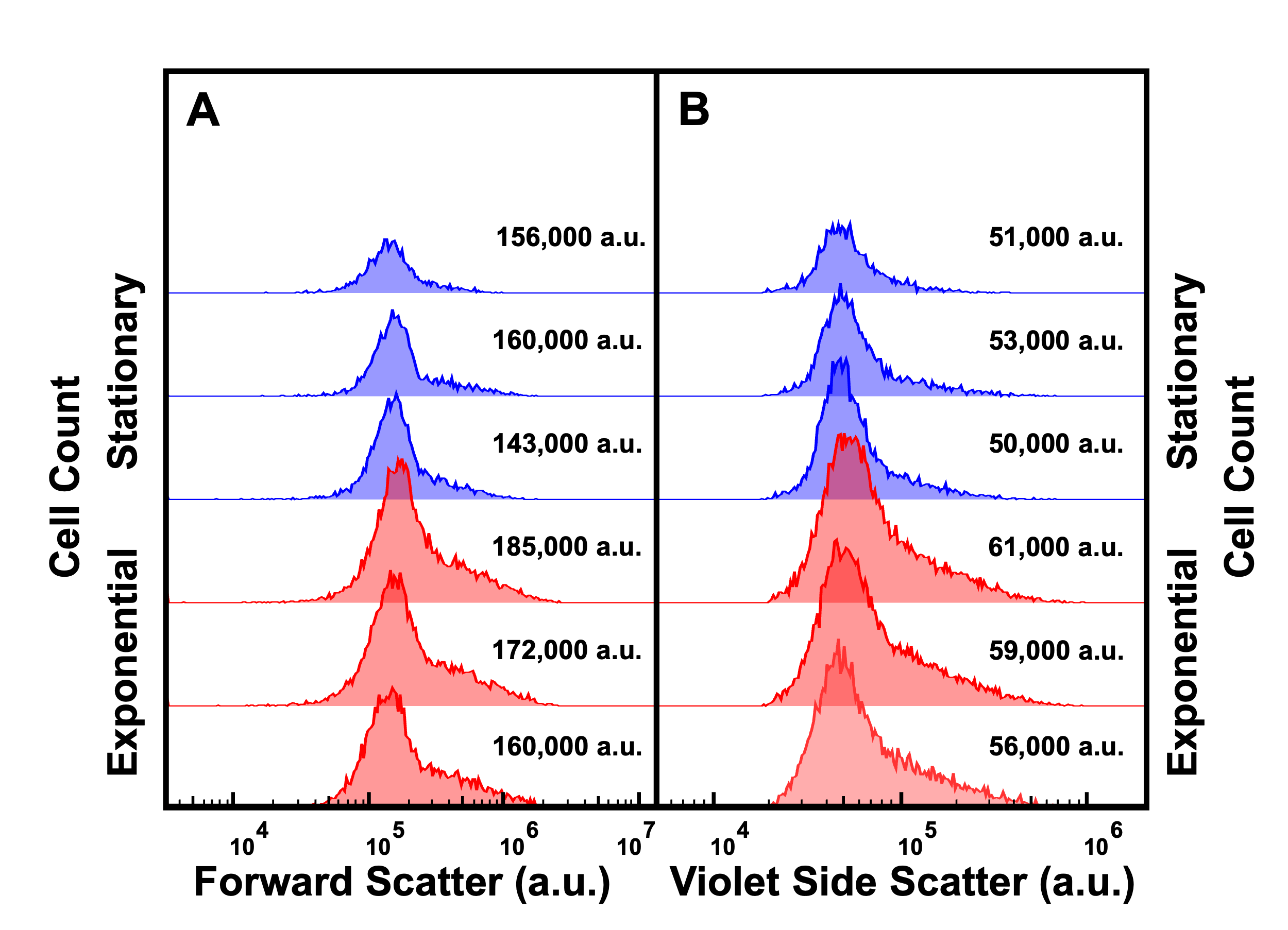

**Figure S17. Impact of the bacterial phase of growth on cell size.** (A and B) Histograms showing the forward scatter (A) and side scatter (B) of high tachyplesin-NBD accumulators in stationary (blue) and exponential (red) phase cells after treatment in 46 μg mL^-1^ (18.2 μM) tachyplesin-NBD in M9 at 37 °C for 60 min with the corresponding median values reported for each distribution. Each distribution is representative of three independent biological replicates.

**Biological processes and genes that are differentially regulated in low compared to high tachyplesin-NBD accumulators**

Low accumulators displayed signs of an overall less translationally and metabolically active state across the entire transcriptome. This is particularly evident in cluster 2 where gene ontology enrichment of differentially expressed genes included processes involved in creating, utilising, and managing cellular components and energy, including protein synthesis, cellular organisation, energy production and gene expression. These genes were found to be downregulated in low accumulators relative to high accumulators. Notably, genes encoding phosphoglucomutase (*pgm*) and translation initiation factor IF-3 (*infC*) were downregulated in low accumulators. Other downregulated genes including the ribosomal proteins (*rpl*, *rpm*, and *rps* operons), RNA polymerase subunits (*rpo* operon), DNA gyrases (*gyr* operon), DNA replication (*parC*, *topA*), and DNA repair (*mutY*, *ruvA*) (see Table S1 and Data Set S3 for a short and complete list of genes involved in these processes, respectively).

In cluster 4, low accumulators displayed an overall enhancement of processes enriched for the transport of organic substances, peptides, dipeptides, and nitrogen compounds, as well as the localisation and assembly of proteins and cell projections, contributing to cellular organisation, population proliferation, and maintenence of cellular homeostasis. These processes involved genes that were predominantly upregulated to a greater extent in low accumulators than high accumulators. Notably, genes encoding MFS efflux pump components such as EmrK, MdtM and MdtL, were upregulated in low accumulators. Several genes encoding ABC transporters were also upregulated in low accumulators, including the *ddp*, *dpp*, and *ydc* operons as well as HisM. Consistent with the data in cluster 4, low accumulators upregulated genes in other clusters encoding the MFS multidrug transporter MdfA and MdtG, putative ABC family antibiotic exporters DppA, OppC, MdlA, YadG, YejA, the Sap operon, the RND multidrug efflux pump membrane fusion protein MdtEF, and the outer membrane protein TolC. It is worth noting that a minority subset of efflux-related genes, i.e. *acrB*, *mdtK*, *mdtA*, *ybhRS* and *yhhJ* were found to be downregulated in low accumulators, suggesting that tachyplesin might not be a substrate of these efflux pumps in *E. coli*. Finally, outer membrane porins OmpC and OmpF were downregulated in low accumulators, potentially reducing the membrane permeability towards tachyplesin (Table S1 and Data Set S3).

Low and high accumulators displayed differential expression of biological pathways underpinning the synthesis and assembly of the lipopolysaccharides (LPS), which is the first site of interaction between gram-negative bacteria and AMPs. Cardiolipin synthase B ClsB, 3-oxoacyl-[acyl-carrier-protein] synthase 1 FabB, LPS-assembly lipoprotein LptE and LPS-export protein LptG were contained in cluster 2 and downregulated in low accumulators. In contrast, genes associated with LPS core biosynthesis (*gmhA*), lipid-A biosynthesis (*lpxA*), and phospholipid biosynthesis (*psd*) belonging to cluster 2 and were found to be upregulated in low accumulators. In cluster 4, ArnF which is involved in L-Ara4N modification of lipid-A was upregulated in low accumulators. Low accumulators further upregulated genes in cluster 1 (for which no biological processes were enriched), including lipid-A L-Ara4N modification genes (*arnBT*), lysophospholipid transporter (*lplT*), LPS-export system proteins (*lptABC*), LPS-assembly protein (*lptD*), phospholipid biosynthesis (*pgpB*, *pssA*), LPS core biosynthesis (*waaCF*), and cell wall remodelling (*yafK*). In contrast, in cluster 3 (for which no biological processes were enriched), genes involved in lipid biosynthesis, including *lpxC*, *pagP*, and pgpAC were downregulated in low accumulators (Table S1 and Data Set S3).

Low and high accumulators displayed differential regulation of biological pathways underpinning the synthesis of outer membrane vesicles (OMVs) that can confer resistance to AMPs and small molecule antibiotics. Specifically, *ompAC*, *degP*, *mcrB*, *tolAB*, and *pal* were downregulated in low accumulators, and their deletion has previously been associated with increased OMV secretion that can confer resistance to AMPs and small-molecule antibiotics. Moreover, *nlpA*, *lysS*, and *waaCF* were upregulated in low accumulators, and their overexpression has been previously associated with enhanced OMV (Table S1 and Data Set S3).

Low accumulators displayed an upregulation of biological pathways, compared to high accumulators, underpinning the synthesis of peptidases and proteases, suggesting a potential mechanism for degrading tachyplesin. In cluster 2, Xaa-Pro dipeptidase PepQ and protease 3 PtrA were found to be upregulated in low accumulators. D-alanyl-D-alanine dipeptidase DdpX and D-alanyl-D-alanine endopeptidase PbpG were found in cluster 4 and upregulated in low accumulators. In cluster 1, low accumulators upregulated signal peptidase I LepB, cytosol non-specific dipeptidase PepD, periplasmic pH-dependent serine endoprotease DegQ, metalloprotease PmbA, and protease 2 PtrB (Table S1 and Data Set S3).

Many of the TFs responsible for the above expression patterns regulate transport systems, indicating a prominent role of transport systems in low tachyplesin-NBD accumulation. Among the top ten TFs displaying higher inferred activity in low accumulators compared to high accumulators, four regulate transport systems, i.e. Nac, EvgA, Cra, and NtrC (Data Set S4).

Furthermore, we found TFs such as NsrR, CRP, ArcA, Fis, and LRP had greater inferred activity in low accumulators. These TFs act to repress genes involved in translation, transcription, DNA repair, and lipid biosynthesis, including components such as translation initiation factor IF-3 InfC; ribosomal proteins in the *rpl* and *rps* operons, and RpmC; RNA polymerase sigma factor RpoS; DNA gyrase GyrAB; DNA-binding protein HupB; DNA topoisomerase TopA; and phosphatidylglycerol synthesis gene PgpC. TF BasR, an activator of the *arn* operon, responsible for lipid-A L-Ara4N modification and TF Nac, an activator of YafK involved in peptidoglycan biosynthesis, displayed greater activity in low accumulators. Moreover, TFs Lrp, Nac, H-NS, and OmpR, which represses DegP, OmpA, Pal, and TolB; TF CsgD, an activator of lipoprotein-28 NlpA, associated with increased OMV secretion, displayed greater activity in low accumulators. Finally, TF NtrC, which is an activator of dipeptidase DdpX was found to have greater activity in low accumulators, and TF LexA, which represses protease 3 PtrA, displayed reduced activity in low accumulators (Data Set S4).

| **Cluster** | **Regulation in Low Tachyplesin-NBD Accumulators** | |
| --- | --- | --- |
|  | **Up** | **Down** |
| **Translation** | | |
| **2** | - | *infC* |
| **Ribosomal Proteins** | | |
| **2** | - | *rplABCDEFKLMNOQRTUWX*, *rpmACDJ*, *rpsCDEFHJKMNOS* |
| **3** | - | *rpIPSVY*, *rpmI*, *rpsABIPQR* |
| **RNA Polymerase and Transcription** | | |
| **2** | - | *rpoBCS* |
| **DNA Replication and Repair** | | |
| **2** | - | *gyrAB*, *hupB*, *mutY*, *parC*, *topA* |
| **3** | - | *mutMT, ruvA* |
| **Glucose Metabolism** | | |
| **2** | - | *pgm* |
| **Persistence Related Genes** | | |
| **1** | *spot*, *dinG* | - |
| **2** | *relA*, *uvrD*, *plsB*, *phoU* | *crp*, *hupB* |
| **3** | - | *dnaJ*, *dnaK*, *lon*, *soxS* |
| **5** | - | *pspABCDEG* |
| **MFS Efflux Pumps** | | |
| **1** | *mdfA*, *mdtG* | - |
| **4** | *emrK*, *mdtML* | - |
| **ABC Transporters** | | |
| **1** | *mdlA*, *sapACDF*, *yadG* | - |
| **3** | *dppA*, *oppC*, *yejA* | *ybhRS*, *yhhJ* |
| **4** | *ddpABCDF*, *dppBCDF*, *hisM, ydcSTUV* | - |
| **RND Efflux Pumps** | | |
| **1** | *mdtEF* | - |
| **2** | - | *acrB*, *mdtA* |
| **MATE-Type Efflux Pumps** | | |
| **2** | - | *mdtK* |
| **Outer Membrane Efflux Proteins** | | |
| **1** | *tolC* | - |
| **Porins** | | |
| **3** | - | *ompCF* |
| **LPS Biosynthesis, Assembly, Export and Modification** | | |
| **1** | *arnBT*, *lplT*, *lptABCD*, *pgpB*, *pssA*, *waaCF*, *yafK* | - |
| **2** | *gmhA*, *lpxA*, *psd* | *clsB*, *lptEG*, *eptA* |
| **3** | - | *lpxC*, *pagP*, *pgpAC* |
| **4** | *arnF* | - |
| **Fatty Acid Biosynthesis** | | |
| **1** | - | *fabA* |
| **2** | - | *fabB* |
| **Outer Membrane Vesicles** | | |
| **1** | *nlpA*, *waaCF* | - |
| **2** | *lysS* | *pal*, *tolAB* |
| **3** | - | *degP*, *mrcB*, *ompAC* |
| **Peptidases** | | |
| **1** | *lepB*, *pepD*, *yafK* | *prlC* |
| **2** | *pepQ* | *map* |
| **3** | - | *ampH*, *dacC*, *lspA*, *mepM*, *pepBT* |
| **4** | *ddpX*, *pbpG* | - |
| **5** | - | - |
| **Proteases** | | |
| **1** | *degQ*, *pmbA*, *ptrB* | - |
| **2** | *ptrA* | *clpA*, *hflC* |
| **3** | - | *degP*, *ftsH*, *loiP*, *lon*, *ompT*, *prc*, *yccA*, *ydgD* |
| **5** | - | *htpX* |
| **Biofilm and Curli Biosynthesis** | | |
| **3** | - | *bssRS*, *tabA* |
| **4** | *csgCDEFG* | - |

**Table S1. Short list of genes involved in biological processes that are differentially regulated in low compared to high tachyplesin-NBD accumulators.** The complete list of genes involved in these processes is reported in Data Set S3.

| **Name** | **Composition** | **Adduct** | **RT** | **Theoretical mass** | **Experimental mass** | **Formula** | **VIP Score** | **∆ppm** |
| --- | --- | --- | --- | --- | --- | --- | --- | --- |
| **Lipids and metabolites upregulated in low tachyplesin-NBD accumulators** | | | | | | | | |
| LPE 20:4 | - | [M+H] | 1.49 | 502.2928 | 502.2925 | C_25_ H_44_ NO_7_ P | 1.46 | -0.59 |
| SM 34:1 | - | [M+H] | 3.40 | 703.5749 | 703.5751 | C_39_ H_79_ N_2_ O_6_ P | 1.30 | 0.28 |
| PE 32:1 | - | [M+H] | 3.20 | 689.4996 | 689.4997 | C_37_ H_72_ NO_8_ P | 1.48 | 0.14 |
| PE 36:5 | - | [M+H] | 3.25 | 738.5069 | 738.5067 | C_41_ H_72_ NO_8_ P | 1.60 | -0.27 |
| PE 38:5 | - | [M+H] | 3.40 | 766.5382 | 766.5380 | C_43_ H_76_ NO_8_ P | 1.56 | -0.26 |
| SM 34:1 | 18:1; 16:0 | [M-H] | 3.8 | 701.5603 | 701.5603 | C_39_ H_79_ N_2_ O_6_ P | 1.97 | 0.28 |
| LPC 18:1 | - | [M-H] | 1.09 | 520.3408 | 520.3406 | C_26_ H_52_ NO_7_ P | 1.23 | -0.38 |
| **Lipids and metabolites upregulated in high tachyplesin-NBD accumulators** | | | | | | | | |
| Cer 32:332**:3** | 14:2; 18:1 | [M+H] | 3.32 | 506.4568 | 506.4567**45** | C_32_ H_59_ NO_3_ | 1.37 | -0.78 |
| PC 36:3 | 18:0; 18:3 | [M+H] | 3.74 | 788.5851 | 788.5849 | C_44_ H_82_ NO_8_ P | 1.42 | -0.25 |
| LPC 22:1 | - | [M+H] | 1.19 | 578.418 | 578.4160 | C_30_ H_60_ NO_7_ P | 1.46 | -3.45 |
| SM 40:2 | 16:1; 24:1 | [M+H] | 3.48 | 785.6531 | 785.6560 | C_45_ H_89_ N_2_ O_6_ P | 1.52 | 3.69 |
| PE 30:0 | 16:0; 14:0 | [M+H] | 3.08 | 664.4912 | 664.4910 | C_35_ H_70_ NO_8_ P | 1.64 | -0.30 |
| PE 31:0 | 16:0; 15:0 | [M+H] | 3.19 | 678.5069 | 678.5067 | C_36_ H_72_ NO_8_ P | 1.71 | -0.29 |
| PE 32:2 | 16:1; 16:1 | [M+H] | 3.21 | 688.4912 | 688.4910 | C_37_ H_70_ NO_8_ P | 1.63 | -0.29 |
| LPC 18:1 | - | [M-H] | 1.09 | 520.3408 | 520.3403 | C_26_ H_52_ NO_7_ P | 1.91 | -0.96 |
| SM 34:1 | 18:1; 16:0 | [M-H] | 3.8 | 701.5603 | 701.5601 | C_39_ H_79_ N_2_ O_6_ P | 1.97 | -0.28 |
| PC 35:0 | - | [M-H] | 2.49 | 774.6018 | 774.6016 | C_43_ H_86_ NO_8_ P | 1.26 | -0.25 |
| PG 38:7 | - | [M-H] | 1.63 | 791.4868 | 791.4861 | C_44_ H_73_ O_10_ P | 1.63 | -0.88 |
| PS 37:7 | 20:5; 17:2 | [M-H] | 5.69 | 790.4664 | 790.4667 | C_43_ H_70_ NO_10_ P | 1.19 | 0.37 |

**Table S2. Lipids and metabolites upregulated in low or high tachyplesin-NBD accumulators compared to untreated bacteria.** Lipid or metabolite name, composition, adduct, retention time (RT), theoretical and experimentally measured mass, formula, variable importance in projection (VIP), and concentration in parts per million (∆ppm) measured via quadrupole time-of-flight liquid chromatography/mass spectrometry.
